# Thermal cycling-hyperthermia in combination with polyphenols, epigallocatechin gallate and chlorogenic acid, exerts synergistic anticancer effect against human pancreatic cancer PANC-1 cells

**DOI:** 10.1101/548552

**Authors:** Chueh-Hsuan Lu, Wei-Ting Chen, Chih-Hsiung Hsieh, Yu-Yi Kuo, Chih-Yu Chao

## Abstract

Hyperthermia (HT) has shown feasibility and potency as an anticancer therapy. Administration of HT in the chemotherapy has previously enhanced the cytotoxicity of drugs against pancreatic cancer. However, the drugs used when conducting these studies are substantially conventional chemotherapeutic agents that may cause unwanted side effects. Additionally, the thermal dosage in the treatment of cancer cells could also probably harm the healthy cells. The purpose of this work was to investigate the potential of the two natural polyphenolic compounds, epigallocatechin gallate (EGCG) and chlorogenic acid (CGA), as heat synergizers in the thermal treatment of the PANC-1 cells. Furthermore, we have introduced a unique strategy entitled the thermal cycling-hyperthermia (TC-HT) that is capable of providing a maximum synergy and minimal side effect with the anticancer compounds. Our results demonstrate that the combination of the TC-HT and the CGA or EGCG markedly exerts the anticancer effect against the PANC-1 cells, while none of the single treatment induced such changes. The synergistic activity was attributed to the cell cycle arrest at the G2/M phase and the induction of the ROS-dependent mitochondria-mediated apoptosis. These findings not only represent the first *in vitro* thermal synergistic study of natural compounds in the treatment of pancreatic cancer, but also highlight the potential of the TC-HT as an alternative strategy in thermal treatment.

## Introduction

Pancreatic cancer is one of the leading causes in cancer death and remains one of the deadliest solid human malignancies worldwide [1]. Patients with pancreatic cancer are commonly diagnosed at the unresectable stage, and in most cases, patients with advanced pancreatic cancer have a poor response to chemotherapy or radiotherapy. In spite of the fact that therapeutic methods have been improved, the prognosis for pancreatic cancer patients still remains poor with a low five-year survival rate [2]. Therefore, there is a need for continued research in novel agents or alternative therapeutic strategies for treating pancreatic cancers, thereby making an improvement for the patients’ quality of life.

Hyperthermia (HT) has emerged as a promising method for treating cancer over the past decades [3]. It is a procedure exposing the tumor tissue to high temperatures that cause cancer cell damage and death. Researches have shown that HT exhibits therapeutic potential against cancer cells through multiple cellular changes, such as protein denaturation and aggregation, inhibition of DNA synthesis, cytoskeleton disruption, and alteration in the calcium homeostasis [4–6]. In addition, HT can directly activate the immune response against the tumors, increase the tumor oxygenation, and improve the drug delivery [7–9]. Although these encouraging results have expanded our understanding of the cytotoxic effects of HT on the cancer cells, in the case of HT as single treatment, it has been shown not to be sufficient to kill cancer cells [10]. To strengthen the effectiveness of HT, several investigations have explored combinations of HT and other cancer therapies, such as radiotherapy and chemotherapy [11]. It has been demonstrated to be effective against various types of cancer, including pancreatic cancer, in that HT enhanced the cytotoxicity of gemcitabine through the inhibition of nuclear factor kappa B (NF-κB) [12–14]. There have also been reports of gemcitabine and other drugs, such as cisplatin and carbonplatin, combined with HT, that demonstrated the clinical efficacy in patients with pancreatic cancer [15, 16]. These data suggest that HT could modify the cytotoxicity of the anticancer drugs, thereby yielding better outcomes in treating pancreatic cancer. However, the drugs used in these combined treatments are conventional chemotherapeutic drugs, which have been known to cause unpleasant and even dangerous side effects.

Nowadays, there has been an increasing interest in natural compounds research due to their lower toxicity and diverse biological properties. Phenolic compounds are among the most studied in cancer prevention and cure, and also the largest group of phytochemicals, as well as being widely distributed in our diet. Particularly, regular intakes of dietary polyphenols have been linked to lower risks of many cancers [17]. Tea and coffee are two of the most consumed beverages worldwide, and the natural phenolic compounds, epigallocatechin gallate (EGCG) and chlorogenic acid (CGA), are the major components in both drinks, respectively. It has been shown that EGCG and CGA have healthy benefits such as antioxidative, anti-inflammatory, and anticancer activities [18–20]. Recently, published findings from animal experimental [21, 22] and clinical studies [23, 24] have demonstrated the ability of EGCG and CGA to suppress tumor cell growth such as breast, lung and bladder cancer. On the other hand, there are a number of studies indicating that natural compounds, including polyphenols, can act as heat synergizers, and thereby improve the anticancer effect [25–29]. Therefore, this has prompted us to conduct the first combined experiment of HT and natural phenolic compounds in pancreatic cancer. Furthermore, we propose a novel approach named thermal cycling-hyperthermia (TC-HT), which allows cells to receive the equivalent thermal dosage through the repeated heat-and-cold cycle. As a matter of fact, there is some evidence indicating that HT lacks tumor selectivity and could also cause harm to normal cells [30, 31]. In this thermal cycled strategy, the cells could avoid prolonged continuous heating under thermal treatment, thereby reducing the toxicity of HT to highlight the synergistic anticancer efficiency of the combined therapy. Therefore, the aim of this study was to investigate the synergistic anti-pancreatic cancer effect of the TC-HT with the CGA and the TC-HT with the EGCG.

In this paper, we examined the effects of the EGCG or CGA combined with the TC-HT on the growth inhibition of PANC-1 cells and evaluated the cell cycle regulation, apoptosis, and the expression of associated proteins to elucidate their underlying mechanisms. Our results demonstrate that the co-administration with the TC-HT and the EGCG or CGA significantly inhibited the cell proliferation and increased the cell death by inducing the cell cycle arrest and mitochondrial apoptotic pathway. These findings first indicate that both the EGCG and CGA might be effective heat synergizers and that the maximum synergistic effects on the PANC-1 cells could be obtained when the TC-HT was administered. Furthermore, the TC-HT as a thermal treatment could be much gentler and feasible, mainly because of the view that the TC-HT itself is relatively harmless to the cells. We believe that this study provides the interesting concept of cyclic thermal application in treatment of cancer *in vitro*, and highlights the potential of the TC-HT as an alternative thermal treatment.

## Materials and methods

### Cell culture

Human pancreatic cancer cell line PANC-1 was obtained from the Bioresource Collection and Research Center of the Food Industry Research and Development Institute (Hsinchu, Taiwan). Cells were plated in 75 cm^3^ cell culture flasks and grown in high-glucose Dulbecco’s modified Eagle’s medium (DMEM) (Hyclone) supplemented with 10% fetal bovine serum (FBS) (Hyclone) and 1% penicillin-streptomycin (Gibco) in a humidified 5% CO_2_ incubator at 37°C.

### Drug treatment and TC-HT exposure

The EGCG was dissolved in distilled water at a concentration of 20 mM, and the CGA was dissolved in dimethyl sulfoxide (DMSO)(Sigma). Subsequently, the two samples were stored at −20°C. The stocks were diluted with a culture medium to the indicated concentration for treatment before usage, and the final concentration of DMSO in each well was 0.05% (v/v). After overnight incubation at 37°C, cells were treated with various concentrations of EGCG or CGA, as well as the solvent vehicle (0.05% DMSO) for CGA. For the TC-HT, the cells were subject to a high and low temperature water bath using a temperature controller modified by a PCR machine (Fig 1A). The cells were exposed to HT (1-cycle TC-HT) at 46°C for 30 min without a break or treated with 46-30°C TC-HT through 3, 6, and 10 cycles of 10, 5, and 3 min, respectively, to receive the equivalent thermal dosages (Fig 1B). In the combined treatment, cells in a medium containing the EGCG or CGA were exposed to the TC-HT. The temperatures of the cancer cells actually sensed were monitored by placing a needle thermocouple at the bottom of the well (Fig 1C). During heat exposure experiment, the non-heating groups (non-treated and drug-treated groups) were exposed to a similar ambient environment as the heating groups over the experimental period of time. After the treatment was completed, cells were stored in a cell culture incubator until the time of analysis.

**Fig 1.**
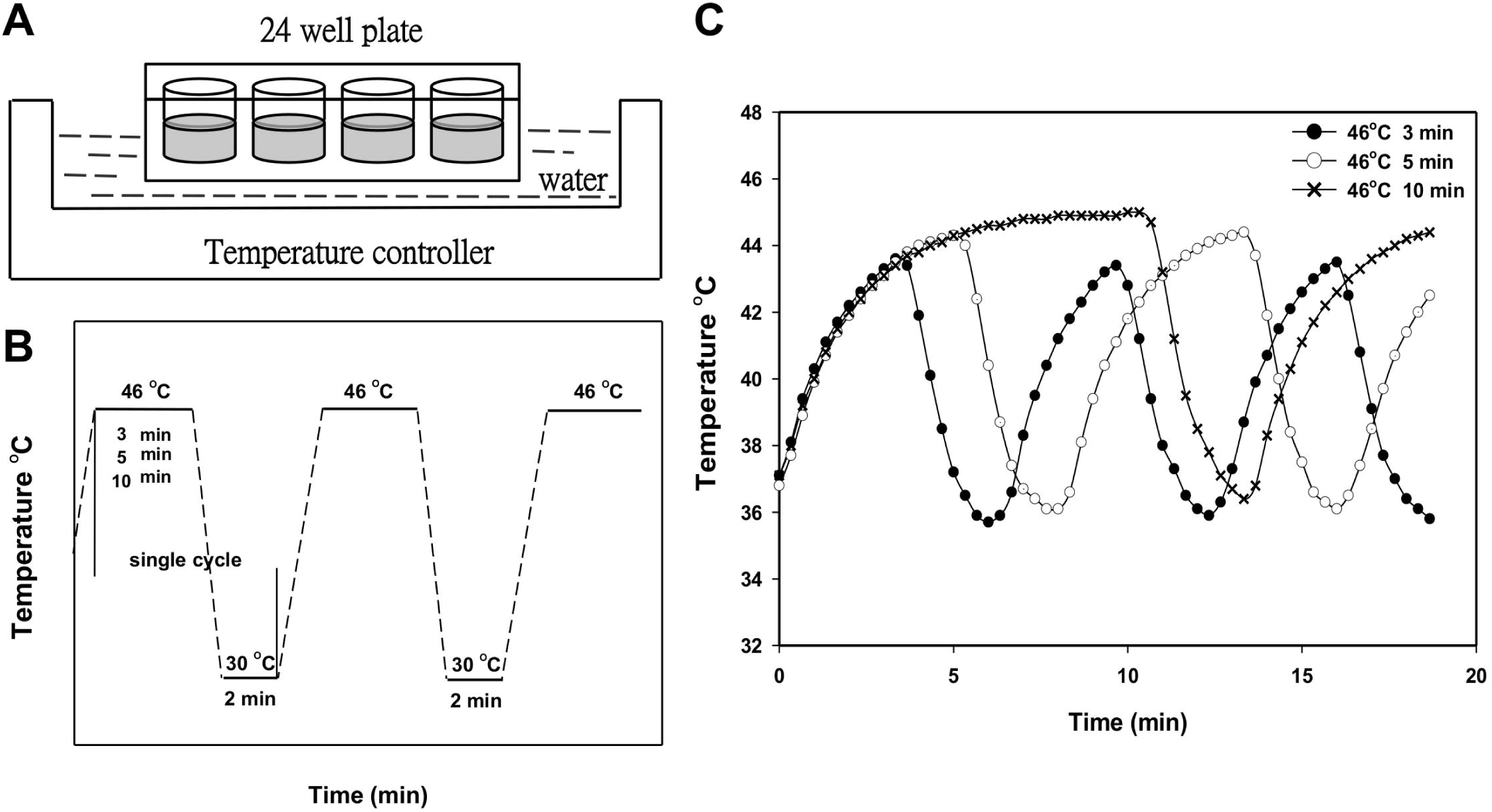
Exposure to the TC-HT via a temperature controller. (A)(B) Schematic of experiment setup and the TC-HT programs setting. (C) The actual temperature recorded every 20 sec in the PANC-1 cells throughout the exposure period.

### MTT assay

Cell viability was accessed by 3-(4,5-dimethylthiazol-2-yl)-2,5-diphenyltetrazolium bromide (MTT) (Sigma) assay. Cells were seeded in 24-well plates and incubated overnight at 37°C. After a single-agent treatment or combined treatment of the TC-HT with the 0.05% DMSO (vehicle control), EGCG or CGA for 24 h, the medium was removed and the cells were washed with phosphate buffered saline (PBS). Cells were then incubated in DMEM containing 0.5 mg/ml MTT for an additional 4 h at 37°C. Then the medium was removed and DMSO was added to dissolve the formazan crystals. The supernatant from each sample was transferred into 96-well plates, and the absorbance was read at 570 nm using an ELISA microplate reader. The calculation of synergism quotient (SQ) was dividing the combined effect by the sum of individual effects. The given treatment shows a synergy when SQ is greater than 1.0.

### Clonogenic survival assay

PANC-1 cells were seeded at 1000 cells/dish in 35 mm Petri dishes for 24 h and treated with 0.05% DMSO (vehicle control), CGA, EGCG and TC-HT alone or in combination. Cell medium was replaced after the treatment, and the dishes were cultured in a humidified 5% CO_2_ incubator at 37°C for additional 14 days. At last, the cells were fixed with 4% paraformaldehyde (PFA) (Sigma) for 10 min and stained with 0.1% crystal violet (Sigma). The colonies containing more than 50 cells were counted, and the number of colonies in each treatment group was normalized to control group.

### Cell cycle analysis

Cells were seeded in 35 mm Petri dishes and incubated overnight at 37°C. After 24 h treatment with DMSO (vehicle control), CGA, EGCG and TC-HT alone or in combination, the cells were harvested, washed with PBS, and fixed with 70% ethanol at 4°C for 30 min. The cells were then stained with propidium iodide (PI) (BD Bioscience) and RNase A (Thermal Scientific) for 30 min in the dark. The stained cells were subject to the cell cycle analysis by using a flow cytometer (FACSCanto II; BD Biosciences), and data were analyzed with ModFit LT software.

### DAPI staining assay

DAPI staining was used to detect morphological characteristics of the nucleus. Cells were cultured on glass coverslips in 35 mm Petri dishes. At the end of each 24 h treatment, cells were washed twice with PBS and fixed with 4% PFA for 10 min at room temperature. After washing twice with PBS, the glass coverslips with attached cells were mounted using Fluoroshield mounting medium with DAPI (Abcam) and examined with a fluorescence microscope (Axio Imager A1; ZEISS).

### Annexin-V/PI double staining assay

Apoptosis was determined by using the Annexin V-FITC/PI apoptosis detection kit (BD Biosciences). Briefly, PANC-1 cells were treated with the TC-HT combined with the DMSO (vehicle control), CGA or EGCG, and then the cells were harvested with trypsin-EDTA (Gibco) and collected by centrifugation at 2,000 × g for 5 min, washed twice with cold PBS, and resuspended in binding buffer containing Annexin V-FITC and PI. The cell suspensions were incubated for 15 min at room temperature in the dark and analyzed by a FACS Calibur flow cytometer.

### Mitochondria membrane potential (MMP) measurement

The cells treated with 0.05% DMSO (vehicle control), CGA, EGCG and TC-HT alone or in combination for 24h were harvested, and resuspended with PBS followed by staining with 20 nM DiOC_6_(3) (Enzo Life Sciences International Inc.) for 30 min at 37°C in the dark. The fraction of cells showing low MMP was then measured by a flow cytometer.

### ROS detection

Cellular Reactive Oxygen Species (ROS) levels of superoxide anion (O_2_^•−^) were detected using the fluorescent dye dihydroethidium (DHE) (Sigma). Cells were treated with the indicated treatments, washed with PBS, and then incubated with 5 μM DHE for 30 min at 37°C in the dark. The fluorescence intensities were measured by flow cytometry and ROS levels were expressed as mean fluorescence intensity (MFI).

### Western blot analysis

After treatment with CGA, EGCG, and TC-HT for 24 h alone or in combination, cells were harvested, washed with cold PBS, and lysed on ice for 30 min in lysis buffer (50 mM Tris-HCl, pH 7.4, 150 mM NaCl, 0.25% deoxycholic acid, 1% NP-40, 1.0% Triton X-100, 0.1% SDS, 1 mM EDTA, 1% phosphate and protease inhibitor cocktail) (Millipore). Cell lysates were clarified by centrifugation at 23,000 × g for 30 min at 4°C, and the protein concentration in the supernatant fraction was quantified using the Bradford protein assay (Bioshop). Proteins were resolved by 10% SDS-PAGE and electrotransferred onto polyvinylidene fluoride membrane (PVDF) (Millipore) in transfer buffer (10 mM CAPS, pH 11.0, 10% methanol). The membranes were blocked with 5% nonfat dry milk/TBST (blocking buffer) for 1 h at room temperature and then incubated overnight at 4°C with diluted primary antibodies in blocking buffer. The specific primary antibodies against Bcl-2, cleaved caspase-8, cleaved caspase-9, cleaved caspase-3 (Cell Signaling), Bax (Santa Cruz), Cdc2, cyclin B1, cleaved PARP and β-actin (GeneTex) were used. After washing with TBST, the membranes were incubated with HRP-conjugated anti-goat (GeneTex) or anti-rabbit (Jackson Immunoresearch) secondary antibody. Chemiluminescence was detected using WesternBright ECL western blotting reagent (Advansta).

### Statistical analysis

The results were presented as mean ± standard deviation (SD). Statistical analysis using one-way analysis of variance (ANOVA) performed with SigmaPlot software. The results were considered to be statistically significant when *p*-values were less than 0.05. Each experiment was done in triplicate.

## Results

### TC-HT in combination with polyphenols synergistically inhibits proliferation of PANC-1 cells

The TC-HT was performed by a modified PCR machine to raise the temperature to a desired level followed by a rapid return to normothermic temperature. The actual temperature in the cells measured by a needle thermocouple could be elevated from 36°C to 43.5, 44.2 and 44.9°C within 3, 5 and 10 min, respectively, and rapidly returned to the physiological temperature (Fig 1C). The effect of the TC-HT and the CGA or EGCG on cell growth was first explored using MTT assay. As shown in Figs 2A and 2B, cells were treated with various concentrations of the CGA or EGCG for 24 h, and both of the compounds only slightly affected the viability of PANC-1 cells. DMSO (0.05%) treated cells with or without the TC-HT did not show a significant difference in viability compared to untreated control cells. Moreover, the viability of PANC-1 cells in response to treatment with the TC-HT decreased in a cycle-dependent manner. When the TC-HT was combined with either of the two compounds, a significant dose-dependent decrease in viability was observed at all four cycling parameters. Although the 1-cycle TC-HT and either the compound cooperatively reduced the proliferation of PANC-1 cells, the heat alone resulted in significant cytotoxicity to the cells. Notably, there was no difference in the cell viability under the condition of the 10-cycles TC-HT when compared to that in the control group, while it still was capable of working synergistically with either the CGA or the EGCG to exert the anti-proliferative activity. The SQ values for the corresponding treatments are as shown in Table 1. This suggests that both the CGA and the EGCG could exhibit a synergistic cytotoxic effect when co-administered with the TC-HT, particularly in the treatment of 10 cycles. We next performed a clonogenic survival assay to confirm the effect of the TC-HT combined with CGA or EGCG on cell proliferation. As shown in Figs 2C and 2D, colony formation in PANC-1 cells was dramatically reduced following both combined treatments. Based on these data, the 10-cycles TC-HT (43.5-36 °C) in combination with the concentration of 200 μM CGA, or the concentration of 20 μM EGCG, was used for all the following experiments.

**Table 1.** Synergy quotient for CGA or EGCG in combination with the TC-HT.

|  |  | TC-HT |  |  |  |
| --- | --- | --- | --- | --- | --- |
|  |  | 1-cycle | 3-cycles | 6-cycles | 10-cycles |
| CGA (μM) | 100 | 0.84 | 0.86 | 1.08 | 1.51 |
|  | 200 | 1.10 | 1.32 | 1.51 | 2.82 |
|  | 300 | 1.11 | 1.34 | 1.38 | 2.40 |
| EGCG (μM) | 10 | 1.18 | 1.11 | 1.27 | 1.95 |
|  | 20 | 1.28 | 1.35 | 1.76 | 3.09 |
|  | 30 | 1.23 | 1.40 | 1.70 | 2.69 |

**Fig 2.**
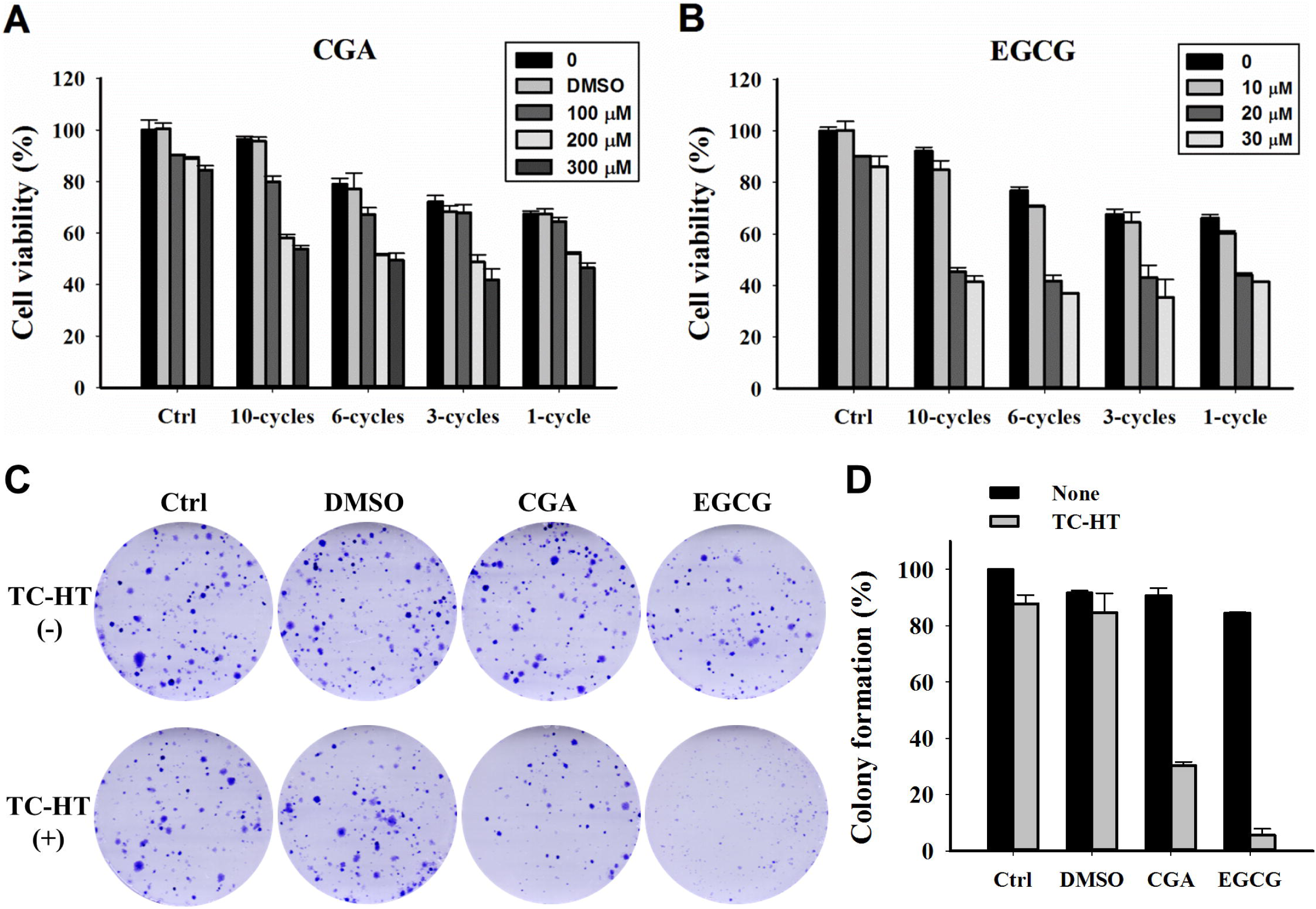
Effects of CGA or EGCG combined with the TC-HT on proliferation in PANC-1 cells. (A) The cells were treated with different cycle numbers of the TC-HT, various concentrations of CGA and 0.05% DMSO (vehicle control) alone or in combination for 24 h. (B) The cells were treated with different cycle numbers of the TC-HT and various concentrations of EGCG alone or in combination for 24 h. (C) Representative images of clonogenic survival assay. (D) Analysis of colony formation rate. Data are presented as mean ± S.D. in triplicate.

### The combination of CGA or EGCG with TC-HT caused G2/M cell cycle arrest in PANC-1 cells

To evaluate the mode of the anti-proliferative effects of the TC-HT, CGA and EGCG, the cell cycle progression in the PANC-1 cells was examined by flow cytometric analysis. As shown in Figs 3A and 3B, treatment with the CGA, EGCG, and TC-HT had no obvious effect on the cell progression when compared with the group of untreated cells. Also, 0.05% DMSO used as the solvent vehicle of CGA either alone or in combination with TC-HT had no effect on the cell progression. Interestingly, the TC-HT combined with either of the two polyphenols resulted in a significant accumulation of cells in the G2/M phase with a concomitant reduction of cells in the G1 phase. To investigate the molecular mechanism of the results of the combined effect on the cell cycle distribution, we examined the relevant proteins involved in the G2/M progression. As shown in Fig 3C, the expression of Cdc2 and cyclin B1 were markedly reduced in response to the combined treatment with TC-HT and CGA or the combined treatment with TC-HT and EGCG. These results indicate that either the CGA or EGCG combined with the TC-HT synergistically induced cell cycle arrest in the G2/M phase by decreasing cyclin B1 and Cdc2 in the cells, and eventually inhibited the cell growth of PANC-1 cells.

**Fig 3.**
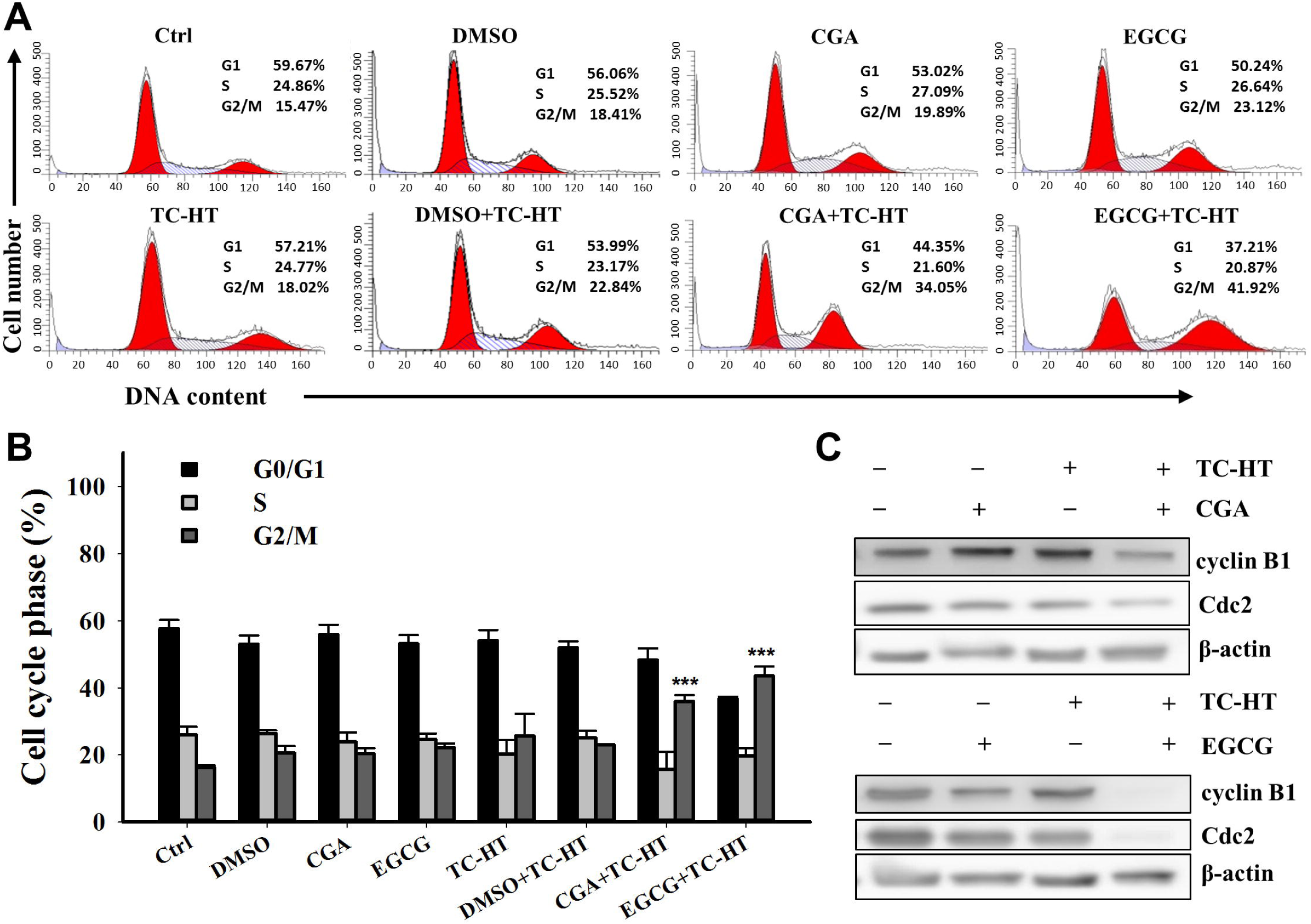
Effects of CGA or EGCG combined with the TC-HT on the G2/M cell cycle arrest in PANC-1 cells. Cells were treated with the TC-HT (10 cycles), 0.05% DMSO (vehicle control), CGA (200 μM), and EGCG (20 μM) alone or in combination (TC-HT + DMSO, TC-HT + CGA, TC-HT + EGCG) for 24 h and stained with propidium iodide (PI) for cell cycle analysis. (A) Representative DNA content profiles. (B) The percentage of cell population in each phase of the cell cycle. (C) Western bolt analysis of the expression of cell cycle regulator proteins cyclin B1 and Cdc2. β-actin was used as an internal control. Data are presented as mean ± S.D. in triplicate. (\*\**p* < 0.01 and \*\*\**p* < 0.001 compared with the untreated control)

### TC-HT combined with CGA or EGCG induces apoptosis in PANC-1 cells

We then investigated as to whether the synergistic cytotoxic effect of the TC-HT with the CGA or EGCG was associated with the induction of apoptosis, and the flow cytometric analysis was performed using Annexin V-FITC/PI staining. As shown in Figs 4A and 4B, the fraction of apoptotic cells after the treatment of PANC-1 cells with either the single agent CGA, or the EGCG or TC-HT were barely increased as compared with untreated control cells. There was no significant difference in apoptosis between the 0.05% DMSO (vehicle control) combined with or without TC-HT and the untreated control. Interestingly, the TC-HT in combination with the CGA significantly increased the early apoptotic cell death in PANC-1 cells (18.6±5.2%). A similar result was also obtained in the combination of the TC-HT and the EGCG, but to a greater extent (24.3±0.8%). These results also showed that the late apoptotic cell death was increased following the exposure of TC-HT with CGA (15.7±4.6%) or EGCG (18.4±4.6%). Furthermore, in contrast to control cells, the DAPI staining demonstrated that the cells treated with either of the two combined treatments had more condensed and fragmented nuclei, which are typical morphological alterations for the apoptosis (Fig 4C). These results suggest that the TC-HT has a potential synergistic effect with the CGA or EGCG on the apoptosis in the PANC-1 cells.

**Fig 4.**
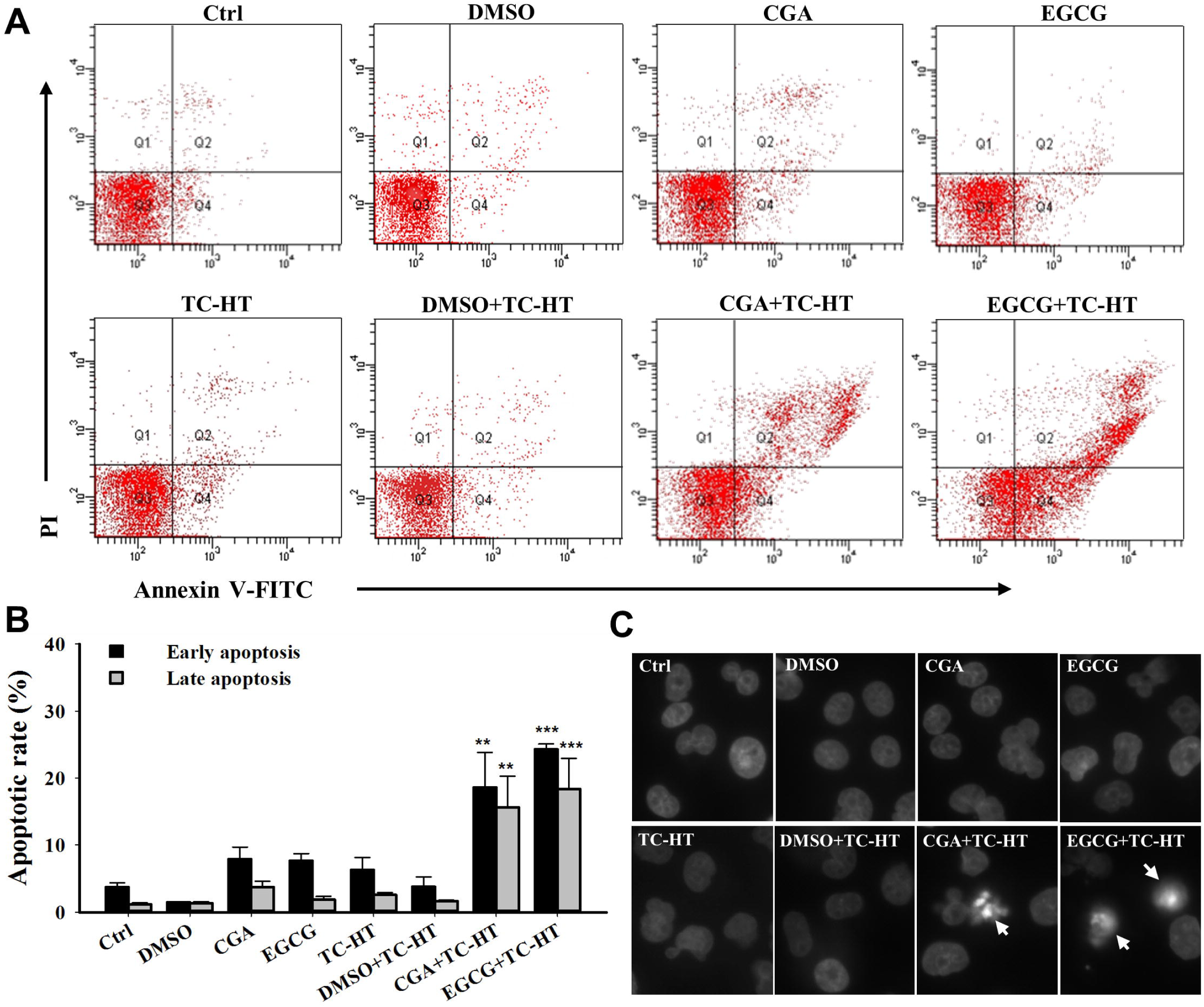
Combination of the TC-HT and the CGA or EGCG induces apoptosis in PANC-1 cells. The apoptosis analysis for the cells following the treatment with the TC-HT (10 cycles), 0.05% DMSO (vehicle control), CGA (200 μM), and EGCG (20 μM) alone or in combination (TC-HT + DMSO, TC-HT + CGA, TC-HT + EGCG) for 24 h. (A) Flow cytometric detection of the apoptosis with Annexin V-FITC/PI double staining. (B) Histogram quantifying the percentage of PANC-1 cells in early and late apoptosis. (C) The nuclei morphology alterations (arrow) were examined using DAPI staining. Data are presented as mean ± S.D. in triplicate. (\*\**p* < 0.01 and \*\*\**p* < 0.001 compared with the untreated control)

### Combination of the TC-HT with CGA or EGCG triggers significant loss in the mitochondrial membrane potential in PANC-1 cells

Collapse of the mitochondrial integrity is a critical event in the cells undergoing apoptosis. To examine whether the combination of the TC-HT and CGA- or EGCG-induced apoptosis involved mitochondrial disruption, the mitochondrial membrane potential (MMP) was assessed using DiOC_6_(3) fluorescence staining by flow cytometric analysis [32]. As shown in Figs 5A and 5B, the treatment with the EGCG, the CGA and its corresponding vehicle control (0.05% DMSO) did not change the MMP level in comparison with the untreated control. In response to treatment with the TC-HT alone or together with vehicle control, there was no significant change in MMP within cells; however, the cells treated with TC-HT showed a stronger effect on the MMP depolarization after the administration of CGA or EGCG (44±6.4% and 75.7±7.6%), which was consistent with the results of the apoptosis analysis as described in Fig 4. These results indicate that the TC-HT combined with the CGA- or EGCG-induced apoptosis in the PANC-1 cells is mediated by mitochondrial dysfunction.

**Fig 5.**
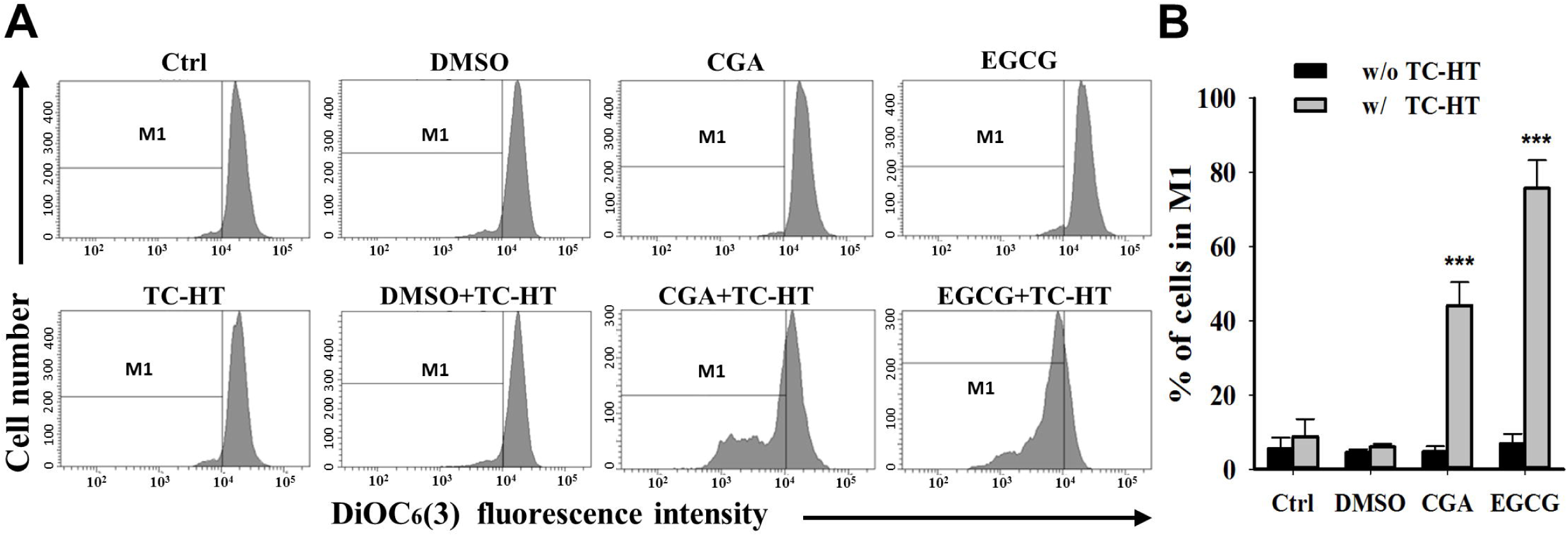
Combination of the TC-HT and the CGA or EGCG induces mitochondrial dysfunction. (A) Flow cytometric analysis of MMP using DiOC_6_(3) after treatment with the TC-HT (10 cycles), 0.05% DMSO (vehicle control), CGA (200 μM), and EGCG (20 μM) alone or in combination (TC-HT + DMSO, TC-HT + CGA, TC-HT + EGCG) for 24 h. The M1 regions indicate the cells with the loss of MMP. (B) Histogram represents the percentage of cells in the M1 region. Data are presented as mean ± S.D. in triplicate. (\*\**p* < 0.01 and \*\*\**p* < 0.001 compared with the untreated control)

### Combination of the TC-HT with CGA or EGCG induces apoptosis through the mitochondrial pathway

It is well known that Bcl-2 family proteins and caspases, along with PARP, play important roles in the mitochondria-mediated apoptosis. To further explore the mechanism by which the TC-HT combined with the CGA or EGCG triggered apoptosis in the PANC-1 cells, we evaluated the expression of the apoptosis-related proteins using western blot analysis. As shown in Fig 6, when compared with control cells, the cleaved caspase-9, −3, and the cleaved PARP were markedly increased after the co-administration with the TC-HT and either the CGA or EGCG. In addition, both of the combined treatments down-regulated the expression of Bcl-2 and up-regulated the expression of Bax, and thus decreased the ratio of Bcl-2 to Bax. These results reveal that the TC-HT combined with the CGA or EGCG promotes apoptosis in the PANC-1 cells via activation of the mitochondrial pathway.

**Fig 6.**
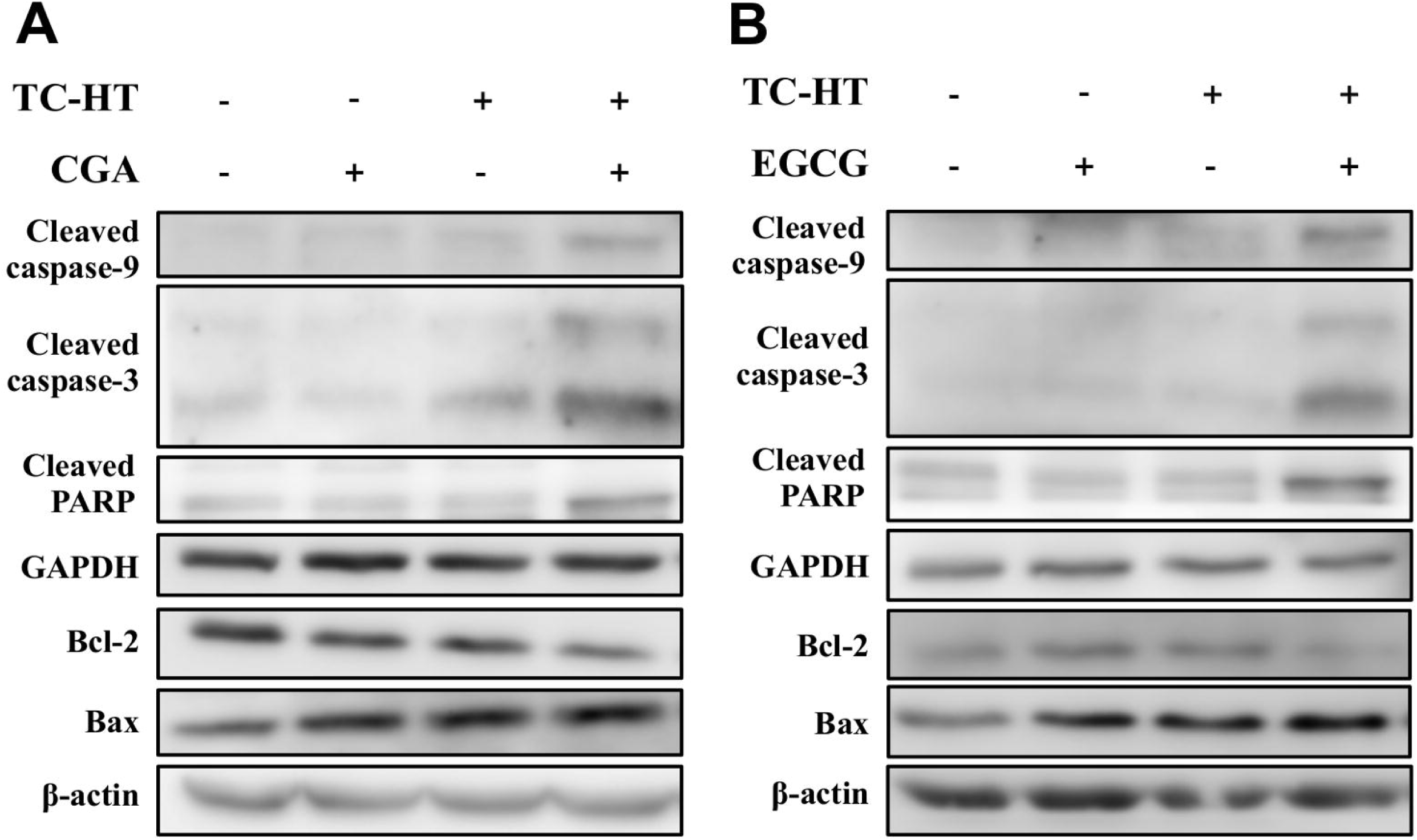
Effects of the TC-HT, CGA and EGCG on the expression of the apoptosis-related proteins in PANC-1 cells. (A)(B) The protein levels of cleaved caspase-9, −3, cleaved PARP, Bcl-2 and Bax of the PANC-1 cells treated with the TC-HT (10 cycles), CGA (200 μM) and EGCG (20 μM) alone or in combination treatment (TC-HT + CGA, TC-HT + EGCG) for 24 h were examined by western blot analysis. GAPDH and β-actin were used as internal controls.

### The role of ROS in the TC-HT combined with the CGA- or EGCG-induced apoptosis in PANC-1 cells

As ROS generation is a critical event in the induction of the apoptosis, we next examined the role of ROS in apoptosis induced by treatment with the TC-HT in combination with either the CGA or EGCG. The intracellular ROS was measured by flow cytometry using a fluorescence probe, DHE, which reacts with O_2_^ −^ [33]. As shown in Fig 7A, the treatment of PANC-1 cells with 0.05% DMSO (vehicle control), CGA or EGCG alone did not alter the level of ROS when compared with the untreated control cells. Also, the level of ROS induced by TC-HT and its combination with DMSO were found to be comparable to the untreated control in the MFI. It is worth mentioning that the TC-HT combined with CGA and the TC-HT combined with EGCG significantly increased the level of ROS in MFI by approximately 2.3- and 3.6-fold, respectively (Fig 7B). These results suggest that the TC-HT combined with the CGA or EGCG may induce apoptosis through the excessive ROS production in the PANC-1 cells.

**Fig 7.**
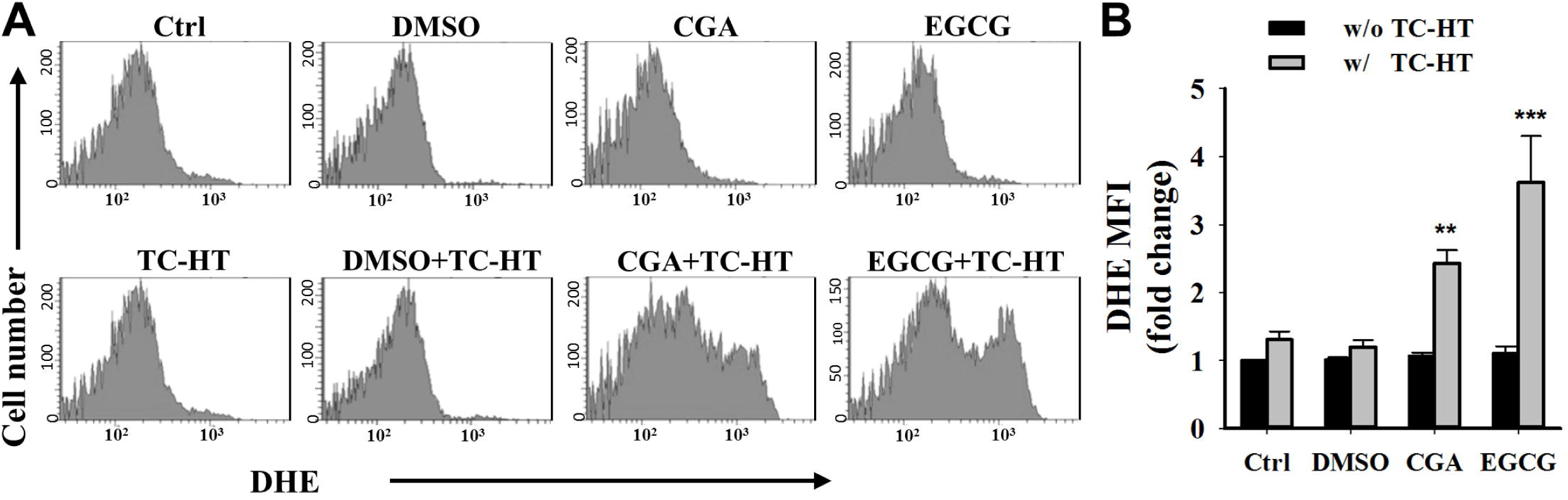
Combined effects of the TC-HT and the CGA or EGCG on ROS generation in PANC-1 cells. (A) The DHE (O_2_^•−^) levels were measured by flow cytometry after treatment with the TC-HT (10 cycles), 0.05% DMSO (vehicle control), CGA (200 μM), and EGCG (20 μM) alone or in combination (TC-HT + DMSO, TC-HT + CGA, TC-HT + EGCG) for 24 h. (B) Graph shows the fold change in MFI of DHE relative to control. Data are presented as mean ± S.D. in triplicate. (\*\**p* < 0.01 and \*\*\**p* < 0.001 compared with the untreated control)

## Discussion

Despite previous studies having shown that HT or polyphenols, the EGCG and CGA, possessed anticancer activities against pancreatic cancer cells [34–37], the effect of these two agents as a combined therapy on pancreatic cancer has not been reported. In this study, we have explored as to whether the herbal compound, CGA or EGCG, could cooperate with heat against the pancreatic cancer cells, and elucidated the cellular mechanism underlying the biological effects. Our results have demonstrated that the exposure to heat using the TC-HT in combination with the CGA or EGCG synergistically inhibited the growth and induced the apoptosis in the PANC-1 cells. Previous studies have shown the benefits of HT as an adjuvant in combination with chemotherapy against many various cancers, including pancreatic cancer [13, 38, 39]. However, there is a major concern that HT can cause unavoidable thermal injury to normal cells. Therefore, it is important to refine the method of heat administration to achieve the desired thermal dosage with the minimal cytotoxicity. To the best of our knowledge, this is the first report indicating a novel approach TC-HT synergizes with the CGA or EGCG on pancreatic cancer PANC-1 cells.

Previous research has reported the response of PANC-1 cells to mild HT at different temperatures [40]. Particularly, the temperature in a range of about 42 to 46°C has been shown capable of inducing the cell death of the pancreatic cell line PANC-1. Our survival analysis revealed a consistent result that the cell growth of PANC-1 in the 1-cycle group was significantly inhibited. Although HT has a cooperative cytotoxicity with the CGA or EGCG, its severe thermal toxicity to a cell is what we want to avoid. The usage of the repeated cycles of heat exposure could provide a means to synergize with the anticancer compounds in the heating process, and avoid the thermal damage accumulation in the following non-heating process. Earlier studies have shown the time-dependent modification of cancer cells during exposure to HT for which the analysis of cellular growth indicated that the survival rate decreased with increasing HT exposure time [41, 42]. Namely, the short exposure of cancer cells to HT may induce cellular stress without affecting their proliferation, while the prolonged exposure may lead to cytotoxicity. It has also been reported that short HT treatment of PANC-1 cells at the temperature 46°C for 5 min had no effect on cell viability [43]. Therefore, we suggest that the 10-cycles TC-HT induced stress rather than damage in cells during each heating cycle. As expected, the cells treated with the TC-HT of different cycling parameters exhibited a decreased cytotoxicity with the increased cycles. It is worth noting that 10 cycles of the TC-HT (43.5-36 °C), or either compound, did not cause a significant alteration in the viability, however, the combination of 10 cycles of the TC-HT with the CGA or EGCG strongly resulted in a dose-dependent decline of the cell viability. Similar results from colony formation were also obtained with the cells treated with the combination of TC-HT and CGA or the combination of TC-HT and EGCG. These results suggested that the either CGA or EGCG may act as heat synergizer and co-administered with the use of the TC-HT could have significantly combined effects against pancreatic cancer PANC-1 cells. Furthermore, we have also tested one liver cancer (HepG2) and one non-cancerous (HEK293) cell lines to examine the combined action. Results shown in S1 Fig revealed that the 10-cycles TC-HT did not cause cytotoxicity in both cancer and non-cancerous cells. Interestingly, such substantial cell death in the PANC-1 cells treated with the TC-HT in combination with CGA or EGCG was not observed in the HepG2 and HEK293 cells, which indicated cell specificity of the combination treatments.

HT has been demonstrated to inhibit the growth of human cancer cells through interfering with the cell cycle progression [27, 29, 44]. In our results, we found that the TC-HT did not cause an obvious accumulation of cells in the G2/M phase. When cells undergoing the TC-HT were given the CGA or EGCG concurrently, the proportion of cells in the G2/M phase was significantly increased. Additionally, this observation was supported by the marked downregulation of Cdc2 and cyclin B1 proteins in the combination of the TC-HT and the CGA or EGCG. The Cdc2 (cell division cycle 2), also known as the CDK1 (cyclin-dependent kinase 1), is a core regulator that drives the cells through G2 and into mitosis. Several studies have reported that the binding of Cdc2 to cyclin B1 complex plays an important role in the G2/M progression [45, 46]. Collectively, these results indicated that the synergistic cytotoxicity of CGA or EGCG under the exposure of TC-HT, at least in part, was associated with the inhibition of the Cdc2/cyclin B1 kinase activity.

Apoptosis, also the best known form of programmed cell death, plays a pivotal role in defending against cancer. The conventional HT has been demonstrated to affect apoptotic pathways in various types of cancer cells [47]. It has also been reported that polyphenols exhibit anticancer activities against different human cancer cells through activating the apoptotic pathway [48, 49]. However, in our study, the results of apoptosis analysis performed using FITC Annexin V and PI double staining indicated that neither the TC-HT nor these two compounds induced significant apoptotic cells (Fig 4). Only when the TC-HT in combination with the CGA or EGCG was conducted, both of the combined treatments significantly elevated the percentage of the early and late apoptotic cells. This finding was further confirmed by the nuclear morphological alterations of the apoptosis in the TC-HT-treated cells in combination with the CGA or EGCG. The apoptotic process can be divided into the death receptor and mitochondrial pathways, and the mitochondrial pathway of cell death is thought to be the major mechanism of apoptosis in mammals. The Bcl-2 family proteins are key regulators of the mitochondrial apoptotic pathway, which comprise of both the anti-apoptotic and pro-apoptotic members. Unbalanced Bcl-2/Bax ratio within the cells induces the disruption of the mitochondrial membrane, release of cytochrome *c*, activation of caspases, and the subsequent cleavage of PARP [50, 51]. Results from Figs 5 and 6 indicated that the combined treatment with the TC-HT and the CGA or EGCG markedly decreased the level of MMP and Bcl-2 expression, and increased Bax expression. We also found that the activation of caspase-9, −3, and the cleavage of PARP were promoted with both of the combined treatments (Fig 6). Collectively, these data indicated that the mitochondria-dependent pathway is involved in the synergistic apoptosis following the combined treatments.

Mitochondria are widely believed to be the main cellular source of ROS. The excessive production of ROS could result in mitochondrial dysfunction, which in turn triggers the apoptosis [52]. We then confirmed if the observation of mitochondrial dysfunction in cells treated with the TC-HT in combination with the CGA or EGCG was promoted by ROS generation. Our result showed that the cellular ROS generation was elevated significantly following the administration of combination of the TC-HT with the CGA or EGCG, indicating that the high levels of ROS production played an important role in the apoptosis induced by the combination of the TC-HT with CGA and the TC-HT with EGCG. HT has been known to act as an adjuvant treatment modality to improve the cytotoxicity of several anticancer agents such as chemotherapeutic drugs [53–55] and herbal compounds [56, 57]. It is thought that HT can enhance the sensitivity of cancer cells to treatment with drugs [58], thereby presenting synergistic anticancer activity. Therefore, these observations of this study tempted us to suggest that the heat stress during temperature cycles may contribute to the potentiation of the CGA or EGCG-triggered cytotoxic responses. Most importantly, the TC-HT could serve as a mild approach to synergize with the natural compound, CGA or EGCG. Furthermore, we also investigate the feasibility of the TC-HT by using standard clinical compound gemcitabine shown to exhibit thermal enhancement of cytotoxicity when combined with standard HT [53, 59]. Similar result as the combination of 1-cycle TC-HT and gemcitabine was observed from analysis of cell viability in the combined treatment with the 10-cycles TC-HT and gemcitabine, whereas the 10-cycles TC-HT alone had no effect on cell viability (S2 Fig). This finding suggests that this approach could be potentially extended to other anticancer agent with heat synergy in cancer research. In practices, the type of thermal therapy is usually associated with the location of tumor. At present time, the TC-HT methodology might have difficulty in applying rapidly cycling heat on centrally located malignancies *in vivo*. Further researches need to be conducted to overcome this problem.

In conclusion, a novel method for producing a desired thermal dosage is proposed in which it applies repeated thermal treatments of short exposure to prevent the toxic effects from prolonged exposure. Our findings have demonstrated the TC-HT shows capability of synergizing with the natural compound, CGA or EGCG, and minimizing the thermal damage resulting from HT. The synergistic activity against the PANC-1 cells was performed primarily via the G2/M arrest and the apoptosis induced through the ROS-mediated mitochondrial pathway leading to an imbalance between Bcl-2 and Bax, activation of caspase-9 and −3, and the cleavage of PARP. This study represents an interesting concept of cyclic thermal application in treatment of cancer *in vitro*, which may potentially demonstrate an alternative to heat treatment. Gaining an optimal combined effect for a variety of cancers via the examination of the relation between the TC-HT parameters and other anticancer drugs is an attractive subject that warrants further investigation.

## Acknowledgments

We would like to thank Technology Commons in College of Life Science, National Taiwan University for use of flow cytometry system, and the staff of the imaging core at the First Core Labs, National Taiwan University Hospital for technical assistance.

## Author Contributions

**Conceptualization:** Chih-Yu Chao.

**Data Curation:** Chueh-Hsuan Lu.

**Formal analysis:** Chueh-Hsuan Lu, Wei-Ting Chen, Chih-Hsiung Hsieh, Yu-Yi Kuo, Chih-Yu Chao.

**Funding acquisition:** Chih-Yu Chao.

**Investigation:** Chueh-Hsuan Lu, Chih-Yu Chao.

**Project Administration:** Chih-Yu Chao.

**Resources:** Chih-Hsiung Hsieh, Chih-Yu Chao.

**Supervision:** Chih-Yu Chao.

**Validation:** Chueh-Hsuan Lu.

**Writing – original draft:** Chueh-Hsuan Lu, Wei-Ting Chen, Chih-Yu Chao.

**Writing – review & editing:** Chih-Yu Chao.

## Submission

***PLOS ONE***

https://journals.plos.org/plosone/

Manuscript received 25 May 2018; 1^st^ revision 31 Oct 2018; 2^nd^ revision 2 Jan 2019; 3^rd^ revision 5 Mar 2019; 4^th^ revision 29 Apr 2019; accepted for publication 16 May 2019.

## Supporting information

**S1 Fig.**
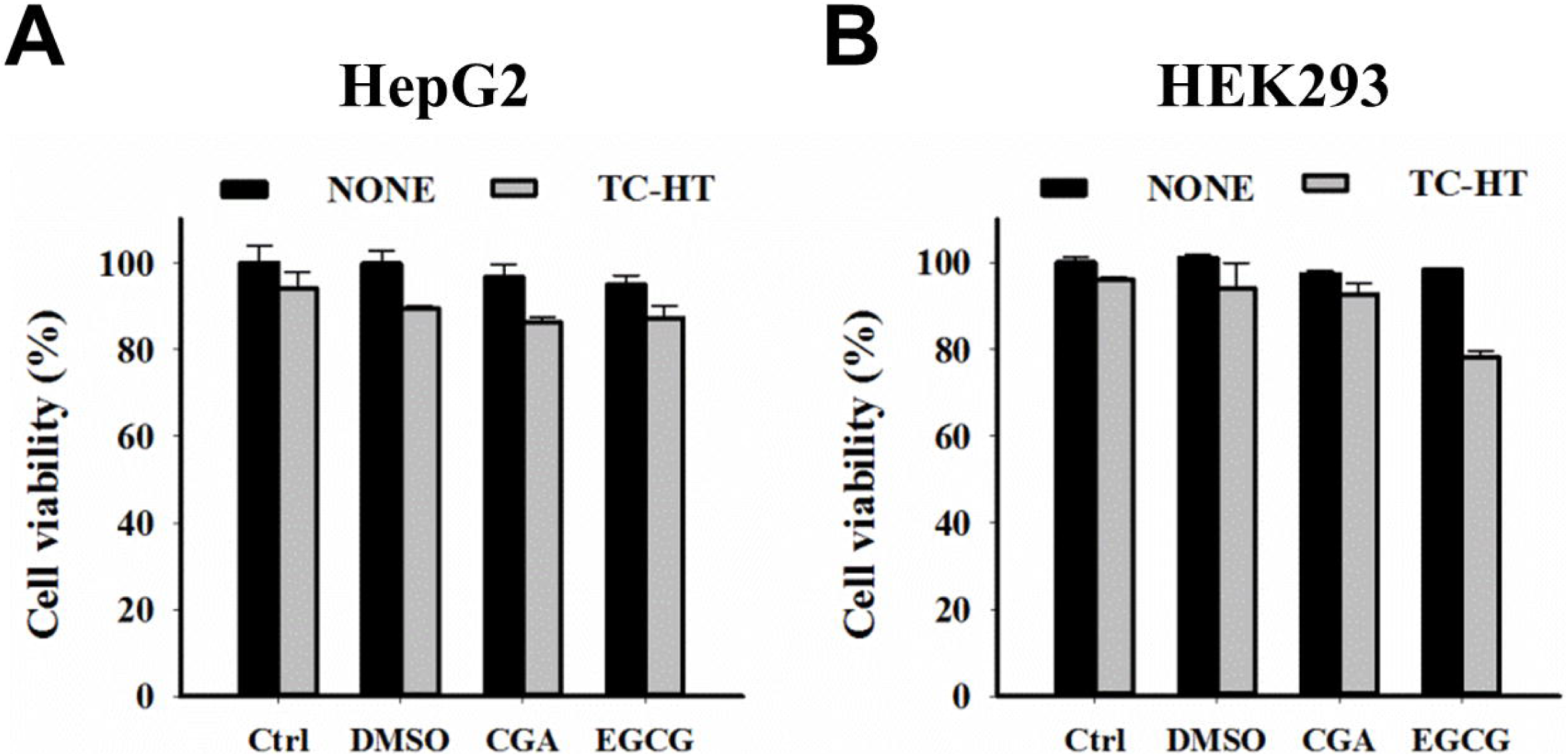
(A) HepG2 and (B) HEK293 cell viability results from MTT assay after the treatments of CGA, EGCG, and TC-HT (10 cycles) alone or in combination (TC-HT + CGA, TC-HT + EGCG) for 24 h.

**S2 Fig.**
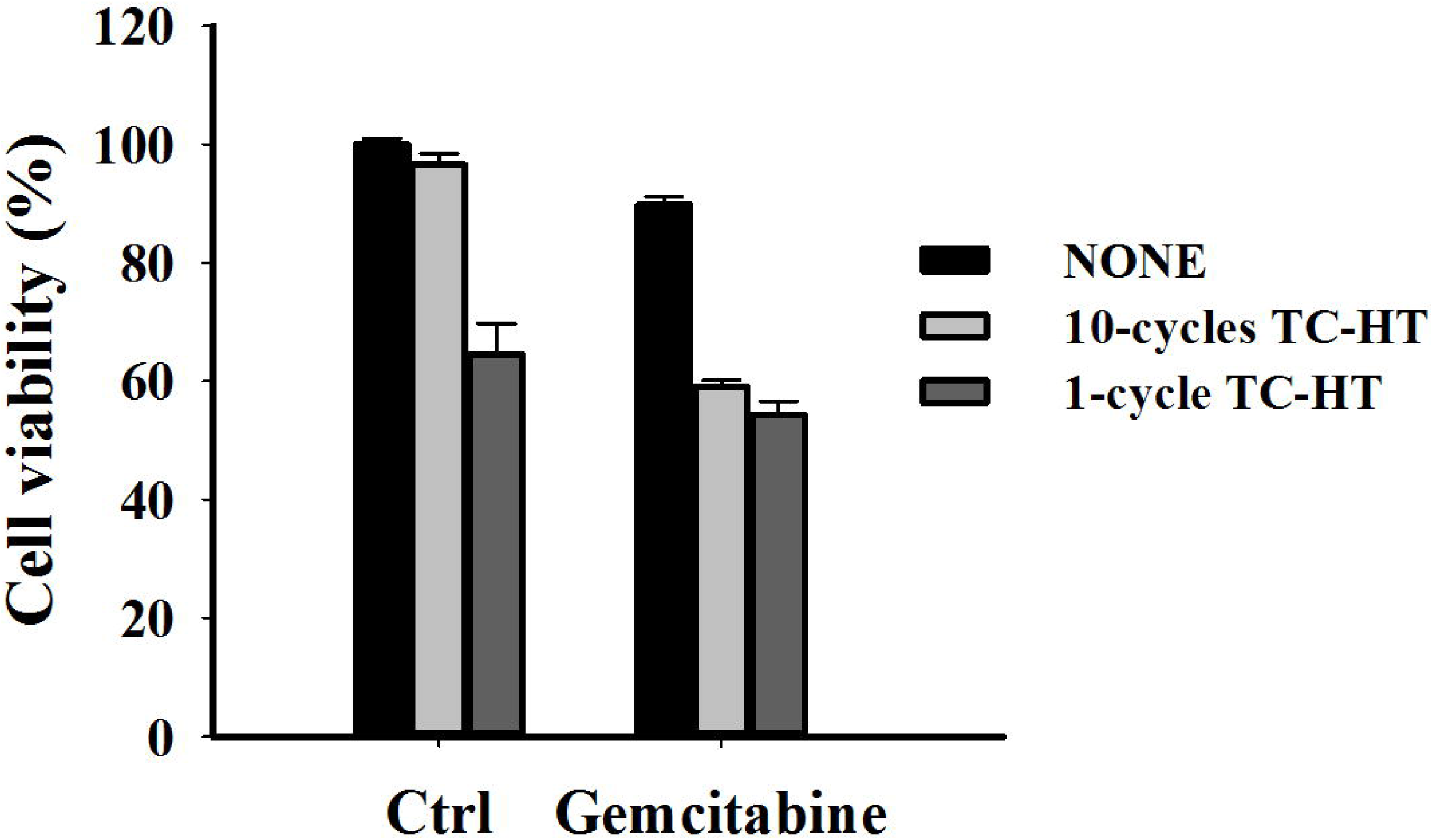
PANC-1 cell viability results from MTT assay after the treatments of gemcitabine (5 μM) and TC-HT (1 cycle and 10 cycles) alone or in combination for 24 h.

**S3 Fig. Raw figures of western blot analysis.**

**S1 File. Raw data of cell viability and colony formation results.**

**S2 File. Raw data of various time points of temperatures.**

**S3 File. Raw data of flow cytometry result.**

